# No evidence for dominance-discovery tradeoffs in *Pheidole* (Hymenoptera: Formicidae) assemblages

**DOI:** 10.1101/2020.05.20.106864

**Authors:** Cristian L. Klunk, Marcio R. Pie

## Abstract

Understanding the mechanisms that allow species coexistence across spatial scales is of great interest to ecologists. Many such proposed mechanisms involve tradeoffs between species in different life-history traits, with distinct tradeoffs being expected to be prevalent at varying temporal and spatial scales. The dominance-discovery tradeoff posits that species differ in their ability to find and use resources quickly, in contraposition to their ability to monopolize those resources, a mechanism analogous to the competition-colonization tradeoff. We investigated the occurrence of this structuring mechanism in *Pheidole* (Hymenoptera: Formicidae) assemblages in Atlantic Forest remnants. According to the dominance-discovery tradeoff, we should observe a consistent interspecific variation along the axis of discovery and dominance. We established 55 sampling units across two sites, with each unit consisting of a sardine bait monitored for three hours. There was no distinction among *Pheidole* species in their ability to find or dominate food sources, suggesting that the dominance-discovery tradeoff does not explain their coexistence. The low levels of aggression between *Pheidole* species could prevent the establishment of dominance hierarchies, whereas the species order of arrival at food sources could allow for resource partitioning through priority effects.

## Introduction

Species coexistence is a fundamental theme in community ecology. How some regions are able to harbor high levels of diversity is still far from being understood, despite a long history of investigation (Hutchinson 1959). Also challenging is to understand how closely related species, which share many aspects of their niches, are able to coexist at fine spatial scales (Hutchinson 1961). Many ecological mechanisms proposed to date to explain species coexistence consider the occurrence of tradeoffs in the performance of species in some life-history traits, which affect their spatial distribution and interspecific interactions (Kneitel and Chase 2004). Species can differ in their competitive and colonization abilities, the levels of abiotic stresses and predation pressures they tolerate, their degree of habitat specialization, among other characteristics, and those differences could explain species coexistence across distinct spatial scales (Kneitel and Chase 2004). Given that diversity translates to ecosystem functioning, the study of species coexistence is essential to understand and manage ecosystem services.

The competition-colonization tradeoff is one of the most classical tradeoff mechanisms proposed to explain species coexistence by suggesting that some species are better at dispersing and colonizing new habitat patches, whereas others are better at displacing incumbent species once they finally colonize the patch (Kneitel and Chase 2004). Although the spatial scale considered under this mechanism can vary, that tradeoff purports to explain coexistence in long temporal scales. Likewise, the competition-colonization tradeoff inspired the proposition of an analogous mechanism that operates in a reduced time range, the dominance-discovery tradeoff (Parr and Gibb 2012). This mechanism proposes that species can differ in their ability to find and dominate resources, and was suggested to explain the coexistence of bees (Hubbell and Johnson 1978; Nagamitsu and Inoue 1997), rodents (Brown et al. 1994) and ants (Fellers 1987; Lebrun and Feener 2007; Feener et al. 2008; Perfecto and Vandermeer 2001; Bertelsmeier et al. 2015*a*).

Ants are important components of terrestrial ecosystems worldwide, particularly in the tropics, where they account for a substantial portion of the animal biomass (Fittkau and Klinge 1973), show considerable species diversity (Economo et al. 2018) and are essential to sustain many ecosystem services (Del Toro et al. 2012). Most ant species are food generalists (Kaspari 2001), because of the marked differences in brood and adult nutritional requirements (Blüthgen and Feldaar 2010). Therefore, if several ant species co-occur and share common resources, one would expect that they might show some level of competition, which suggests the existence of mechanisms of coexistence to sustain high levels of diversity. However, the role of competition in ant community structure is far from settled (Parr and Gibb 2010), despite having traditionally been considered as a “hallmark” of ant communities (Hölldobler and Wilson 1990). Competition in ant communities can be manifested in two ways. Through interference, some ant species exclude competitors from resources by direct aggression or territoriality, whereas by exploitative behaviors some species are specialized in finding and removing resources before other species arrive (Davidson 1998). This leads to dominance hierarchies, in which some species (dominants) are capable of monopolizing resources, whereas others are better at finding resources quickly and removing them before the arrival of the dominants (subordinates) (Fellers 1987; Feener et al. 2008; Arnan et al. 2012; Yitbarek and Philpott 2019).

Ant species are considered as dominant, in the context of those hierarchies, either by the outcomes of pairwise interactions with other species, where the winners are considered dominants (behavioral dominance), or by a large recruitment of workers on food sources (numerical dominance), preventing the use of the resource by other species (Davidson 1998). The dominance-discovery tradeoff is an interesting mechanism to explain coexistence between subordinate and dominant ant species. To find resources at faster rates, an ant colony needs to optimize the number of scout workers (Pearce-Duvet and Feener 2010) that search for food, as opposed to dominant species that invest more in recruiters (Fellers 1987). This tradeoff was tested in several situations, including in intraspecific interactions (Jordan and Blüthgen 2007), between invasive species (Bertelsmeier et al. 2015*a*) and within guilds (Fellers 1987). Although some studies demonstrated the importance of this mechanism in warranting ant species coexistence (Fellers 1987; Lebrun and Feener 2007; Feener et al. 2008; Bertelsmeier et al. 2015*a*), others questioned its ubiquity (Adler et al. 2007; Jordan and Blüthgen 2007; Parr and Gibb 2012; Stuble et al. 2013; Castracani et al. 2014; Camarota et al. 2018; Antoniazzi et al. 2021; Dáttilo and MacGregor-Fors 2021). Some authors also stressed factors that can modify the outcome of dominance-discovery tradeoffs (Davidson 1998; Feener 2000; Adler et al. 2007; Parr and Gibb 2012).

The genus *Pheidole* is one of the most diverse ant lineages, with more than 1000 species described to date (Bolton 2021), showing high levels of diversity in the subtropical and tropical regions (Economo et al. 2015; 2019). The combination of high levels of local diversity and the fact that most *Pheidole* species are food generalists and consequently share many resources (Wilson 2003), raise the question of what mechanisms could explain such high levels of species co-occurrence. Here we test, through bait experiments, the possibility that *Pheidole* species co-exist at local spatial scales by sharing resources according to the predictions of the dominance-discovery tradeoff. If the dominance-discovery tradeoff explains the coexistence of *Pheidole* species, we expect to observe a consistent interspecific variation along the axis of discovery (order of arrival) and dominance (number of workers recruited) (Fellers 1987). Alternatively, a lack of evidence for the dominance-discovery tradeoff could be the result of *Pheidole* species being more spatially segregated than the expected by chance (Connor and Simberloff 1979; Gotelli 2000), so they would rarely interact directly to dispute resources. We therefore test whether *Pheidole* species are spatially segregated, and expect that they are not more segregated than expected by chance.

## Material and Methods

We carried out field experiments in two neighboring urban forest fragments located in the campus of the Universidade Federal do Paraná in Curitiba, Paraná, southern Brazil (Site A - approximately 5.5 ha; 25°26′45.9″S; 49°13′55.5″W and site B - approximately 15 ha; 25°26′52.06″S; 49°14′19.24″W). The vegetation in these sites is classified as mixed ombrophylous forest (Reginato et al. 2008), which is part of the Atlantic Forest biome (IBGE 2012). The climate in the study region is subtropical humid with temperate summers, categorized as *Cfb* according to Köppen’s system (Alvares et al. 2013). Experiments were carried out between the end of February and the beginning of April of 2018, during the warm season, always in the morning, to minimize the effects of climatic differences between sampling days. We established 55 sampling units along three transects in each of the two sampling sites (25 and 30 in sites A and B, respectively). In each transect, sampling units were set 10 meters apart from each other to avoid resampling individuals from the same colony in different sampling units (Bestelmeyer et al. 2000). To properly follow the behavior of ant workers at each sampling unit during each experiment, only five sampling units were established each day, for a total of 11 sampling dates over the course of the study. Each sampling unit consisted of a bait including 2-mL mixture of sardine and wheat flour, placed on a Petri dish (9 cm in diameter) that prevented ants from having access to the bait from below, which could hamper direct observation and potentially underestimate their presence on the baits. We offered a large amount of sardine to avoid quick removal of the resource before the end of the experimentation time. There was no evidence that Petri dishes prevented workers to access the sardine. *Pheidole* species are heavily attracted to sardine baits, which represent a rich source of proteins, sodium and lipids, and it has been demonstrated that protein resources are more attractive to ants in forest remains at earlier successional stages (Bihn et al. 2008). Each experiment lasted for three hours, such that each sampling unit was checked every 20 min, for a total of nine time intervals, and the same person led all experiments.

We estimated the number of major and minor workers of *Pheidole* species at the sampling units with the aid of a manual counter, and recorded the occurrence of negative interactions with other *Pheidole* species and with non-*Pheidole* species (defined as interactions where a direct contact between workers leads them to get away from each other, without necessarily a physical injury). In the case of interactions between *Pheidole* species, the same interaction was counted twice, once for each species, given that we were not always able to define which species led the attack against the other. For later species identification, we collected one individual per subcaste and morphospecies at each sampling unit and deposited them in micro tubes filled with 95% ethanol. Voucher specimens were deposited at the “Coleção Entomológica Padre Jesus Santiago Moure”, Zoology Department, UFPR. Identifications followed the *Pheidole* taxonomic key available in Wilson (2003). No permits were required for fieldwork, and collections of specimens followed Brazilian legal provisions.

### Analysis

If species were spatially segregated due to competitive interactions (Connor and Simberloff 1979; Gotelli 2000), we would rarely see more than one *Pheidole* species in the same sampling unit, which would necessarily lead to a pattern in which the species that discovered the bait will remain alone until the end of the experiment. To test if species are more spatially segregated than expected by chance, we calculated the C-score (Stone and Roberts 1990) based on presence-absence matrices considering all sampling units. The observed C-score was then tested for significance against an estimated C-score obtained from null models. We generated 5000 random matrices with fixed row and fixed column sums (Gotelli 2000). We estimate the C-score for the combined data as well as the data for each sampling site independently and for the common species (defined as the species that occurred in more than one sampling unit). The C-score analysis were carried out in EcoSim7.0 (Gotelli and Entsminger 2001).

To assess species discovery ability, we defined a discovery ability index (DAI), which ranges from 0 to 1 and is defined as the number of sampling units a species arrived first, divided by the total number of sampling units that the species occurred at. If two or more *Pheidole* species occurred together in the same sampling unit at the first time interval, both were treated as discoverers. Although observations started after 20 minutes, which prevent us from ascertaining the exact order of arrival in a few instances in which two *Pheidole* species were presented in the first time interval (four sampling units), there was no evidence that this time was enough for a species to discover the bait and be completely removed by another *Pheidole* species, a process that tends to demand a greater amount of time (Perfecto and Vandermeer 2011). To determine dominance ability as a proxy for numerical dominance (Davidson 1998), we defined a dominance index (DI), which also ranges from 0 to 1 as the number of sampling units that a *Pheidole* species dominated, divided by the total number of sampling units in which that species occurred. A species dominated the sampling unit if it recruited more minor workers at the final time interval than the median number of minors recruited across all baits and time intervals in which it occurred (Table 1), which resembles the monopolization index proposed in Castracani et al. (2014) and Antoniazzi et al. (2021), based on the original proposition of Santini et al. (2007). In considering the median number of minors recruited by each species to the baits as a threshold to define numerical dominance, instead of a pre-defined specific number of workers for all species, we avoid the issues associated with interspecific differences in recruitment behavior.

**Table 1.** Species sampled, approximate number (median) of major and minor workers registered, occurrence at baits, number (frequency) of negative interactions among *Pheidole* species and between *Pheidole* species (Neg. int. *Pheidole*) and non-*Pheidole* species (Neg. int. non-*Pheidole*). Species occurrence represents the number of sampling units that each species occurred at out of the total number of sampling units (55). The frequency of negative interactions among *Pheidole* species is defined as the number of time intervals that each species engages in a negative interaction with another *Pheidole* species out of the total number of time intervals that it co-occurred with any other *Pheidole* species. The frequency of negative interactions between *Pheidole* species and non-*Pheidole* species is defined as the number of time intervals that a *Pheidole* species engages in a negative interaction with a non-*Pheidole* species out of the total number of time intervals that it occurred.

| Species | N° major records<sup>a</sup> | N° minor records<sup>a</sup> | Occurrence (%) | Neg. int. *Pheidole* | Neg. int. non-*Pheidole* |
| --- | --- | --- | --- | --- | --- |
| *P. aff. bucculenta* | 0 (0) | 4 (1) | 1.82 | 1(25) | 0(0) |
| *P. aff. longiseta* | 24 (2) | 345 (30) | 3.64 | 2(25) | 2(15.38) |
| *P. aff. rosae* | 0 (0) | 209 (4.5) | 7.27 | 10(52.63) | 2(10) |
| *P. aff. susannae* | 211 (25) | 900 (100) | 1.82 | 0(0) | 0(0) |
| *P. aff. tijucana* | 3 (0) | 109 (10) | 1.82 | 1(20) | 0(0) |
| *P. cf. lucretii* | 218 (2) | 3833 (60) | 18.18 | 6(28.57) | 14(20) |
| *P. flavens* complex | 15 (0.10) | 1544 (5) | 45.45 | 19(20.43) | 9(5.81) |
| *P. hetschkoi* | 1 (0) | 699 (4) | 32.73 | 14(36.84) | 15(14.29) |
| *P. nesiota* | 35 (1) | 197 (6) | 7.27 | 2(40) | 1(5) |
| *P. risii* | 0 (0) | 831 (5) | 20 | 5(17.24) | 0(0) |
| *P. sp.1* | 4 (0) | 335 (40) | 1.82 | 1(14.29) | 0(0) |
| *P. sp.2* | 0 (0) | 16 (2) | 1.82 | 0(0) | 2(28.57) |

<sup>a</sup>Approximate sum (median) of major and minor workers at each sampling unit at each time interval. This sum does not correspond to abundance, since it is likely that the same individuals were present in the same Petri dish over the entire experiment time. If more than 100 workers were counted, we stopped to count and considered this as the final number of workers.

We calculated species occurrence, the frequency of negative interactions among *Pheidole* species and between *Pheidole* and non-*Pheidole* species. The details of these calculations are provided in Fig. 2, Tables 1 and S3. To test if the occurrence of negative interactions influences the number of workers being recruited to the sampling units, we estimated the median number of minor workers being recruited at sampling units where negative interactions occurred among *Pheidole* species and between *Pheidole* and non-*Pheidole* species. We applied a paired Wilcoxon test (Zar 2010*a*) to compare if these numbers are different from the median number of minors recruited at all sampling units, which is the number considered to define the DI. To test if the dominance-discovery tradeoff influences the coexistence of *Pheidole* species, we applied a Spearman rank correlation test (Zar 2010*b*) between the DI and DAI, with a negative correlation being considered as support for the dominance-discovery tradeoff (Fellers 1987). We combined the data of both study areas but also considered them independently, as well as the data from only the common species. Analyses were carried out in R 4.0-2 (R Core Team 2020).

**Fig. 1.**
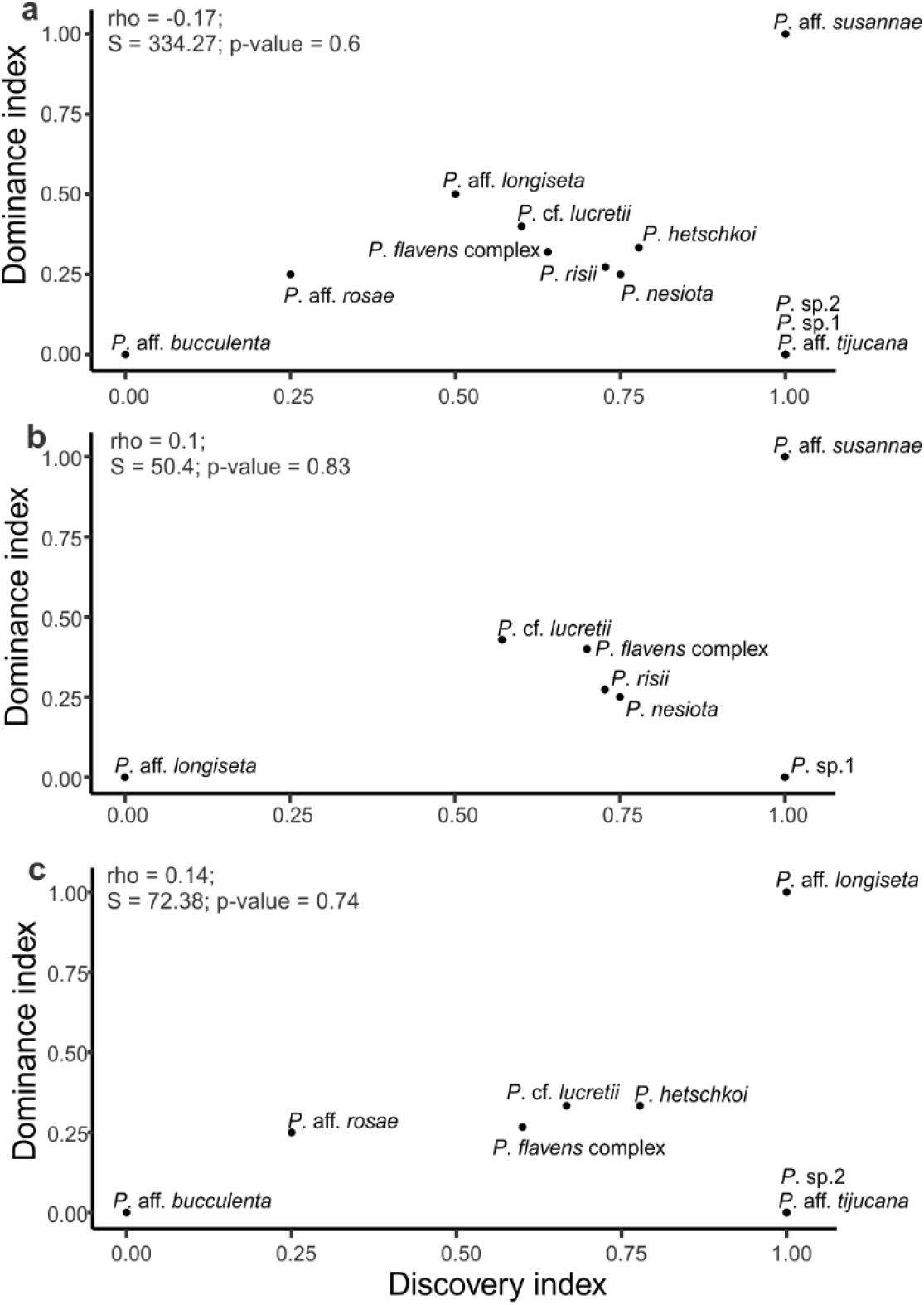
Correlations between discovery and dominance indices from the combined (a), site A (b) and site B (c) data. Insets display the results of the Spearman correlation tests. Plots were generated with the R package GGPLOT2 (Wickham 2009).

**Fig. 2.**
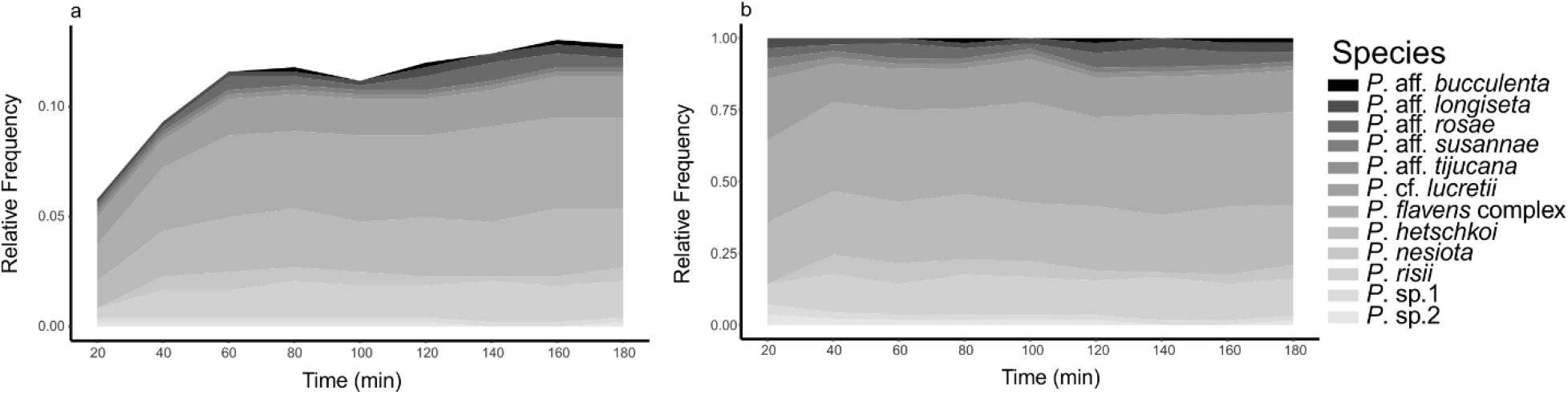
Relative frequency of *Pheidole* species over the experiment time (a). Relative frequency considers the number of records of each *Pheidole* species out of the total number of time intervals that a *Pheidole* species occurred (483). In (b), the same relationship is visualized with the relative frequencies of each species standardized to a constant height. Plots were generated with the R package “ggplot2” (Wickham 2009) and “viridis” (Garnier 2018).

## Results

We recorded 12 *Pheidole* species across our two sampling sites (Table 1). Two morphospecies were indistinguishable in the field, so to avoid any under or overestimation of each morphospecies worker numbers we combined their data under a unique species cluster, the *P. flavens* complex. *Pheidole* species visited 48 out of 55 sampling units during our field experiments. Of all sampling units, 47% were discovered by only one species, 24% by two species and 16% by three species. The most frequent species was *P*. *flavens* complex (45.45%), followed by *P*. *hetschkoi* Emery, 1896 (32.73%) and *P. risii* Forel, 1892 (20%) (Table 1). *Pheidole* aff. *susannae, P*. cf. *lucretii* and *P*. sp. 1 recruited the highest number of minor workers, whereas *P*. aff. *bucculenta* and *P*. sp. 2 recruited the lowest numbers (Table 1). Four species never recruited major workers (*P*. aff. *bucculenta*, *P*. aff. *rosae*, *P*. *risii* and *P*. sp. 2), and only *P*. aff. *susannae* and *P*. cf. *lucretii* recruited more than five majors at any sampling unit and time interval (Table 1).

Regarding negative interactions, *Pheidole* sp.2 did not exhibit negative interactions against other *Pheidole* species and *P*. aff. *susannae* never co-occurred with another *Pheidole* species (Table 1). The remaining ten species showed aggressive behaviors against other *Pheidole* species (Table S1), with *P*. aff. *rosae* showing negative interactions in 52.63% of the time intervals in which it occurred, followed by *P. nesiota* Wilson, 2003 (40%) and *P*. *hetschkoi* (36.84%). However, aggressive behaviors between *Pheidole* species were observed in only 26.07% of time intervals in which at least two *Pheidole* species co-occurred. During the experiments, some non-*Pheidole* species (*Brachymyrmex* sp., *Camponotus rufipes* (Fabricius, 1775), *Gnamptogenys* sp., *Linepithema* sp., *Nylanderia* sp.,*Odontomachus chelifer* (Latreille, 1802), *Pachycondyla striata* Smith, 1858, and *Solenopsis* sp.) were recorded and showed agonistic interactions with *Pheidole* species (Table 1). *Pheidole* sp. 2 interacted negatively against a non-*Pheidole* species in 28.57%of time intervals where it was recorded, followed by *P*. cf. *lucretii* (20%) and *P*. aff. *longiseta* (15.38%). Negative interactions had no effect on the number of minor workers being recruited at sampling units, given that the median number of minors of each *Pheidole* species along the entire experimentation time was not different from the median number of minors being recruited at baits that registered negative interactions among *Pheidole* species (V = 4; p = 0.42) and where negative interactions happened between *Pheidole* and non-*Pheidole* species (V = 7.5; p-value = 0.46). Consequently, there was no evidence that negative interactions influenced the DI.

By comparing the spatial distribution of *Pheidole* species with random distributions generated by null models, we did not find any evidence that species are more aggregated than expected by chance (Table S2; observed ≤ expected, p = 0.314) or less aggregated than expected by chance (Table S2; observed ≥ expected, p = 0.694) for the combined data. When looking at site A individually the same patterns were found (Table S2; observed ≤ expected, p = 0.106; observed ≥ expected, p = 0.919), but the results from site B suggest otherwise that *Pheidole* species are spatially segregated (Table S2; observed ≤ expected, p = 0.967; observed ≥ expected, p = 0.038). However, when we considered only the most common species, the co-occurrence patterns of all sites are not different from what is expected by chance (Table S2). We did not find any negative correlation between DI and DAI (Figure 1), considering both the combined data (Figure 1a) and the independent data from sites A (Figure 1b) and B (Figure 1c). The conclusions remain the same when we considered just the common species (Figure S1). In general, most *Pheidole* species stay on the sampling units throughout the entire duration of the experiment (Table S3; Figure 2). Even rarely recorded species, like *P*. aff. *tijucana* and *P*. aff. *susannae*, which occurred just at one sampling unit, were never replaced for those baits (Table S3).

## Discussion

Despite the relatively high number of co-occurring *Pheidole* species at our sampling sites, we did not find any evidence for dominance-discovery tradeoffs promoting species coexistence, suggesting that *Pheidole* species show similar strategies to invest in resource discovery and dominance. Specifically, we found no negative correlation between the discovery and dominance indices, contrary to the expectation for the occurrence of the tradeoff (Fellers 1987). By reviewing the literature on dominance-discovery tradeoff, some authors suggested that the tradeoff might be context dependent (Davidson 1998; Parr and Gibb 2012), as also proposed under a simulation approach (Adler et al. 2007). The levels of aggressiveness between pairwise species interactions can contribute to establish dominance hierarchies (Savolainen and Vepsäläinen 1988). However, here we recorded negative interactions between *Pheidole* species in only 26.07% of the encounters between at least two *Pheidole* species, and we found no evidence that *Pheidole* species changed their recruitment behavior due to the occurrence of any kind of negative interaction. Although commonly recorded in ant interactions, aggressiveness need not be the norm in interspecific encounters, even among dominant and highly invasive ants (Bertelsmeier et al. 2015*b*). Neutral interactions between coexisting species at bait experiments represent a great portion of the observed outcomes (Blüthgen et al. 2004; Stuble et al. 2017; Gray et al. 2018), yet they are mainly neglected, with emphasis in studies granted preferentially to true agonistic behaviors (Stuble et al. 2017). In *Pheidole* species, the increased proportion of major workers recruited to food sources could be an important trait to enhance the probability of resource dominance (Mertl et al. 2010). Although the species considered here were markedly distinct with respect to the number of majors recruited at baits, this difference was not enough to be translated into a dominance hierarchy, and majors were rarely observed engaging in negative interactions.

Reacting aggressively or not against a potential competitor is context-dependent (Tanner and Adler 2009; Barbieri et al. 2013). The “dear enemy” phenomenon posits that animals can recognize potential competitors that live close to their own territory from others that are strangers, and react more aggressively against strangers (Langen et al. 2000; Tanner and Adler 2009). If the *Pheidole* species that approach the sampling units are neighbors that constantly interact, this could explain the relative low levels of aggression in interspecific encounters. However, the opposite trend was also recorded for ants (Gordon 1989), and to evaluate the importance of this phenomenon in the present context would require a more detailed investigation of the nature of the interspecific encounters between the *Pheidole* species. The type of resource being disputed and the nutritional demands of the colony can also influence the foraging behavior of ant workers (Cornelius and Grace 1997; Kay 2004; Silberman et al. 2016). In general, proteins and lipids are the main nutrients to feed larvae and the queens, whereas for adult workers carbohydrates are the main energy sources to sustain their activities (Blüthgen and Feldhaar 2010). Although we used here a unique source of food to attract *Pheidole* workers to baits, sardine baits are sources of both proteins and lipids (Lasmar et al. 2021), which are commonly searched by *Pheidole* species in the field along with carbohydrate (Rosumek 2017). Further studies can investigate if other food types can elicit more aggressive behaviors between *Pheidole* workers, which occasionally can result in interactions that follow the predictions of the dominance-discovery tradeoff.

Coupled with the low number of agonistic interactions between ant species at baits, in nearly half of our sampling units we recorded only one *Pheidole* species throughout the entire experimentation time. In some situations, it was clear that the first species that arrived at baits had nests close to the baits (author’s pers. obs.), which represents a clear advantage to find the resource and recruit workers to dominate it. The implication is that the crucial stage for resource assurance could be the discovery event, a fact recently suggested for Neotropical arboreal ant communities (Camarota et al. 2018; Antoniazzi et al. 2021) and a Neotropical urban gradient (Dátillo and MacGregor-Fors), which was defined as a discovery-defense strategy (Camarota et al. 2018). Although we did not find a positive relationship between discovery and dominance abilities, there is a trend towards it in our results from both the combined data and individual data from sites A and B, which was not confirmed due to the contrasting behavior of three species (*P*. sp.1, *P*. sp.2 and *P*. aff. *tijucana*). This pattern differs from the two step view whereby the resource used by ants is partitioned between a food search phase, followed by a latter moment of competition displacement and species turnover where other ants have greater success, the so called dominance-discovery tradeoff (Fellers 1987). This suggests that other mechanisms can be involved in structuring this community, such as priority effects (Andersen 2008). Priority effects predict that the order of species arrival at empty patches determines the subsequent structuring of the community (Fukami 2015; De Meester et al. 2016). The order of arrival into a resource patch can determine the level of aggressiveness of dominant ant species (Barbieri et al. 2013), in a way that influences resource partitioning. Order of arrival can also change the colony level of activity and survival (Barbieri et al. 2013), with important consequences for species coexistence and community structure.

By arriving first at baits, the *Pheidole* species in our study have more time to recruit nestmates and dominate the resource before the arrival of other *Pheidole* species, and in the absence of successful takeovers in resource patches, coexistence can be improved (Adler et al. 2007). If the *Pheidole* species investigated here are spatially segregated to some degree, this could explain why so many sampling units were visited by only one species. Although our analysis of spatial structure from site B suggests that those *Pheidole* species are spatially segregated, it seems that this result is an effect of the rare species. When just common species are considered, the patterns of species co-occurrence agree with what is expected by chance (Stone and Roberts 1990; Gotelli 2000). Another possibility is that the complexity of the habitat delays the time to discover and recruit nestmates to the baits. More complex habitats affect the way that ants sense the environment and influence its locomotion capacity (Kaspari and Weiser 1999), which consequently affects the time to discover and recruit nestmates to the baits (Parr and Gibb 2012). The time to discover food sources can vary because of several biological aspects of ant species, but there are cues that in more simple vegetation types ants are able to find resources at faster rates (Gibb and Hochuli 2004) than in more complex habitats (Holway 1999). Although our sampling sites conserve vegetation elements of secondary stages of succession (Reginato et al. 2008), both sites are composed of arboreal species that provide a complex leaf-litter in which ants forage, which can hinder ant locomotion and navigation.

We show here that a classical mechanism of species coexistence, the dominance-discovery tradeoff (Fellers 1987), was not able to explain *Pheidole* species co-occurrence in two assemblages of Atlantic Forest remains in south Brazil. Similarly, coexistence of *Pheidole* species seems not to be related to tradeoffs between dominance and environmental resistance (Tschá and Pie 2019), which suggests that evolutionary tradeoffs can have a secondary role to structuring the *Pheidole* assemblages investigated. Although there are cues for the importance of priority effects (Andersen 2008) to the resource partitioning between *Pheidole* species, this hypothesis remains to be tested in the present community. Also important to advance our knowledge about *Pheidole* species coexistence is a deeper understanding of their dietary preferences (Rosumek 2017; Rosumek et al. 2018) and the role of stochastic mechanisms in promoting species coexistence (Andersen et al. 2013; Stuble et al. 2017). The importance of the dominance-discovery tradeoff in structuring ant communities is highly context-dependent (Davidson 1998; Parr and Gibb 2012), and the prevalence of ecological tradeoffs in general depend on the spatial scales being considered (Kneitel and Chase 2004). Therefore, a lack of evidence for the occurrence of the dominance-discovery tradeoff in many ant assemblages (Parr and Gibb 2012) should not be viewed as a definitive proof against the importance of such mechanisms in the structuring of ant communities. The dominance-discovery tradeoff should be more broadly explored, along different spatial scales and with different organisms, to provide a deeper understanding about the range of their influence in ecological communities to ensure species coexistence.

## Acknowledgments

The authors thank Alexandre Casadei Ferreira for assistance in species identification and Félix Rosumek for comments on the manuscript.

## Authors’ contributions

CLK and MRP designed the study, CLK carried out field and lab work and data analysis, CLK and MRP wrote the manuscript.

